# Use of waggle dance information in honey bees is linked to gene expression in the antennae, but not in the brain

**DOI:** 10.1101/2020.12.07.414078

**Authors:** Anissa Kennedy, Tianfei Peng, Simone M. Glaser, Melissa Linn, Susanne Foitzik, Christoph Grüter

## Abstract

Communication is essential for social animals, but deciding how to utilize information provided by conspecifics is a complex process that depends on environmental and intrinsic factors. Honey bees use a unique form of communication, the waggle dance, to inform nestmates about the location of food sources. However, as in many other animals, experienced individuals often ignore this social information and prefer to rely on prior experiences, i.e. private information. The neurosensory factors that drive the decision to use social information are not yet understood. Here we test whether the decision to use social dance information or private information is linked to gene expression differences in different parts of the nervous system. We trained bees to collect food from sugar water feeders and observed whether they utilize social or private information when exposed to dances for a new food source. We performed transcriptome analysis of four brain parts critical for cognition: the subesophageal ganglion, the central brain, the mushroom bodies, and the antennal lobes but, unexpectedly, detected no differences between social or private information users. In contrast, we found 413 differentially expressed genes in the antennae, suggesting that variation in sensory perception mediate the decision to use social information. Social information users were characterized by the upregulation of dopamine and serotonin genes while private information users upregualted several genes coding for odor perception. These results highlight that decision making in honey bees might also depend on peripheral processes of perception rather than higher-order brain centers of information integration.

## Introduction

Exchanging information is essential in all animal societies. Communicating about resources, reproductive state, group membership, and threats are vital in ensuring the survival and success of the group. However, relying on social information is often not the only available option, e.g. to find a food source, but searching for a resource individually can often be the better choice (Laland, 2004; Kendal et al., 2009; Hoppitt & Laland, 2013; Dechaume-Moncharmont et al., 2005; I’Anson Price et al., 2019). Furthermore, an individual can rely on private information (e.g. spatial memory) about previously visited food source locations (Rendell et al., 2010; Grüter & Leadbeater, 2014). It is crucial for an organism to assess the different available options and their consequences to make the best decision in a given environment. Acquiring information through individual exploration, for instance, provides up-to-date information, but comes with the cost of trial-and-error learning. Social information avoids the costs of individual learning and exploration, but can involve the inefficient or erroneous transmission of information (Giraldeau et al., 2002; Dechaume-Moncharmont et al., 2005; Rieucau & Giraldeau, 2011; I’Anson Price et al., 2019). Thus, animals often employ flexible strategies for deciding between social or private information (Laland, 2004; Kendal et al., 2009; Hoppitt & Laland, 2013; Grüter & Leadbeater, 2014).

Social insects employ various methods to send signals to nestmates. Information exchange regarding resources is particularly well-studied and a wide range of communication behaviors are used, such as tandem running in ants (Alleman et al., 2019, Möglich et al., 1974; Glaser and Grüter, 2018) and trail pheromones in ants, and stingless bees (Jarau, 2009; Hölldobler & Wilson, 2009; Czaczkes et al., 2015). Honey bees (Apini) use a unique form of communication, the waggle dance that gives spatial information to nestmates about both distance and direction of a food source or a nest site in relation to the sun (von Frisch, 1967). In foraging, dances are performed by returning foragers as an advertisement for high quality food sources. Furthermore, waggle dancers emit floral odors and a blend of hydrocarbons that provide additional information and stimulate foraging in unemployed foragers (Gilley et al., 2018; Thom et al., 2007; Farina et al., 2012). Only a relatively small percentage of waggle dance followers use dance information to discover new food sources. The majority of waggle dances trigger experienced foragers to resume foraging at already familiar food sources, disregarding social dance information for private spatial information (Biesmeijer & Seeley, 2005; Grüter et al., 2008). While various factors, like experience (Richter & Waddington, 1993; Biesmeijer & Seeley, 2005; Grüter & Ratnieks, 2011;) and age (Tofilski, 2009; Woyciechowski & Moroń, 2009) are likely to affect whether a bee uses social information, still little is known about the neuronal basis of dance communication and its use (Barron & Plath, 2017).

Numerous studies have shown that social insect behavior and responses to social information are linked to brain gene expression (Toth et al., 2010; Robinson et al., 2008; Zayed & Robinson, 2012; Ingram et al., 2011; Toth & Robinson, 2009). For example, foragers have a unique pattern of gene expression compared to nurses as they upregulate genes associated with synaptic plasticity and cognition (i.e spatial learning and memory), whereas nurses upregulate genes associated with intracellular signaling involved in the transition from nurse to forager (Whitfield et al., 2003). Even among foragers different gene expression patterns can be found. For example, pollen and nectar foragers differentially upregulate genes associated with regulating food intake (Brockmann et al., 2008). Behavioral variation within foragers seems to be strongly connected to the expression of genes that are important in biogenic amine signaling, such as dopamine, octopamine, tyramine, glutamate, and serotonin signaling (Liang et al., 2012; Scheiner et al., 2002; Schulz et al., 2003; Barron et al., 2002; Scheiner et al., 2017a). Indeed, manipulation of biogenic amine levels can alter foraging behavior (Liang et al., 2012; Peng et al., 2020; Linn et al., 2020) and perception of food rewards and odors (Mercer & Menzel, 1982; Barron et al., 2002; Scheiner et al., 2002). Most studies have focused on whole brains to reveal expression differences between behavioral groups (e.g. Whitfield et al., 2003; Liang et al., 2012; Alleman et al., 2019). However, different brain parts serve specific functions and are expected to differ in gene expression. For example, the antennal lobes receive input from the olfactory sensory neurons in the antennae (Paoli et al., 2016) and process olfactory information (Homberg et al., 1989; MaBouDi et al., 2017; Paoli et al., 2016). Insect mushroom bodies are key brain areas for multimodal sensory integration, learning and memory (Strausfeld et al., 2009; Collett & Collett 2018), whereas the central brain supports foraging behavior via motor control (Hanesch et al., 1989). Barron and Plath (2017) have suggested that the central brain might play a crucial role in the decoding of waggle dance information. Finally, the subesophageal ganglion mediates reward and taste perception (Kreissl et al., 1994; Dacks et al., 2005; Sinakevitch et al., 2005).

If and how these different brain parts are involved in dance communication and information-use is not well understood. Furthermore, we still know little about the role of the peripheral nervous system for decision making and information processing (see e.g. Ozaki et al., 2005). The antennae, in particular, play important functions in social insect behavior, both within and outside the colony, such as mediating pheromone signaling (Nagari & Bloch, 2012; Vergoz et al., 2009; Grozinger et al., 2003; Pankiw, 2004), nestmate recognition (Ozaki et al., 2005; van Zweden & D’Ettorre, 2010) and odor learning (Robertson et al., 2006; Rogers & Vallortigara, 2008). An important role of the antennae in mediating different behaviors is also likely to explain why foragers and nurses show distinct antennal expression of chemical sensory and biogenic amine genes (Nie et al., 2018; McQuillan et al., 2012). Chemical stimuli differentiation and odor perception are not only important for task differentiation (Arenas & Farina, 2012; Balbuena & Farina, 2020), but could play a role in the decision to use social or private information (Thom et al., 2007).

Here we compared the gene expression of bees that used dance information (social information, SI) with those that preferred private information (PI) in different brain areas and the antennae in the honey bee *Apis mellifera.* We trained cohorts of workers to sucrose solution feeders and, subsequently, confronted them with conflicting social information about a new high-quality food source. As was shown for scouts, *i.e.* foragers that search for new food sources independently (Liang et al., 2012), we predicted that there are distinct neurogenomic signatures underlying the decision to use either social or private information. We compared different brain and peripheral chemosensory areas in both types of bees. We demonstrate that bees that decode and use waggle dance information differ in gene expression only in the antennae and provide evidence for roles of biogenic amine signaling and olfactory perception.

## Materials and Methods

### Colony Set-up

A total of six observation hives of *Apis mellifera carnica* were studied from August through October 2016 (H1 – H3) and 2018 (C1 - C3), each containing approximately 2000-3000 workers of mixed ages. Colonies were established from the Johannes Gutenberg University apiary in Mainz, Germany, a few weeks prior to the start of experiments. Each of the observation colonies contained three frames, brood, food reserves and were headed by a naturally mated queen.

### Training

Training was conducted one colony at a time. Workers were trained according to standard training procedures to collect sucrose solution at one of two artificial feeders (von Frisch, 1967; Linn et al., 2020). First, a cohort of 50-60 workers was trained to the training feeder (T.F.). These workers were used as the samples that would later be designated as either social or private information users on test day. Then, a smaller cohort of ~20 foragers was trained to the dance feeder (D.F.). These workers would be designated as dancers on test day. Both feeders were 150m from the observation colonies with ca. 160 meters separating the training and dance feeder (see Linn et al., 2020, their Fig. 1). Workers were trained to their respective feeder with an unscented 0.8M sucrose solution and were individually marked with a number tag on the thorax. This spatial arrangement ensured that workers would visit only one feeder and no mixing of individuals between dance and training feeders occurred. The day after training, the sucrose solution was reduced to 0.3M at both feeders with the addition of an identical scent (5μL of essential oil /100mL sucrose solution). This concentration made sure that trained foragers would return to their respective feeder, but not recruit additional bees. Colonies were trained to a different odor: C1, H1 = sage, C2, H2 = jasmine, C3, H3 = peppermint. During 60 minutes, workers were allowed to visit their feeder repeatedly (2016: 5.24 ± 3.79 visits, N = 191; 2018: 8.09 ± 5.17 visits, N = 102). The 60-minute training with scented solution allowed workers to associate reward, scent, and location of the respective feeder.

**Figure 1:**
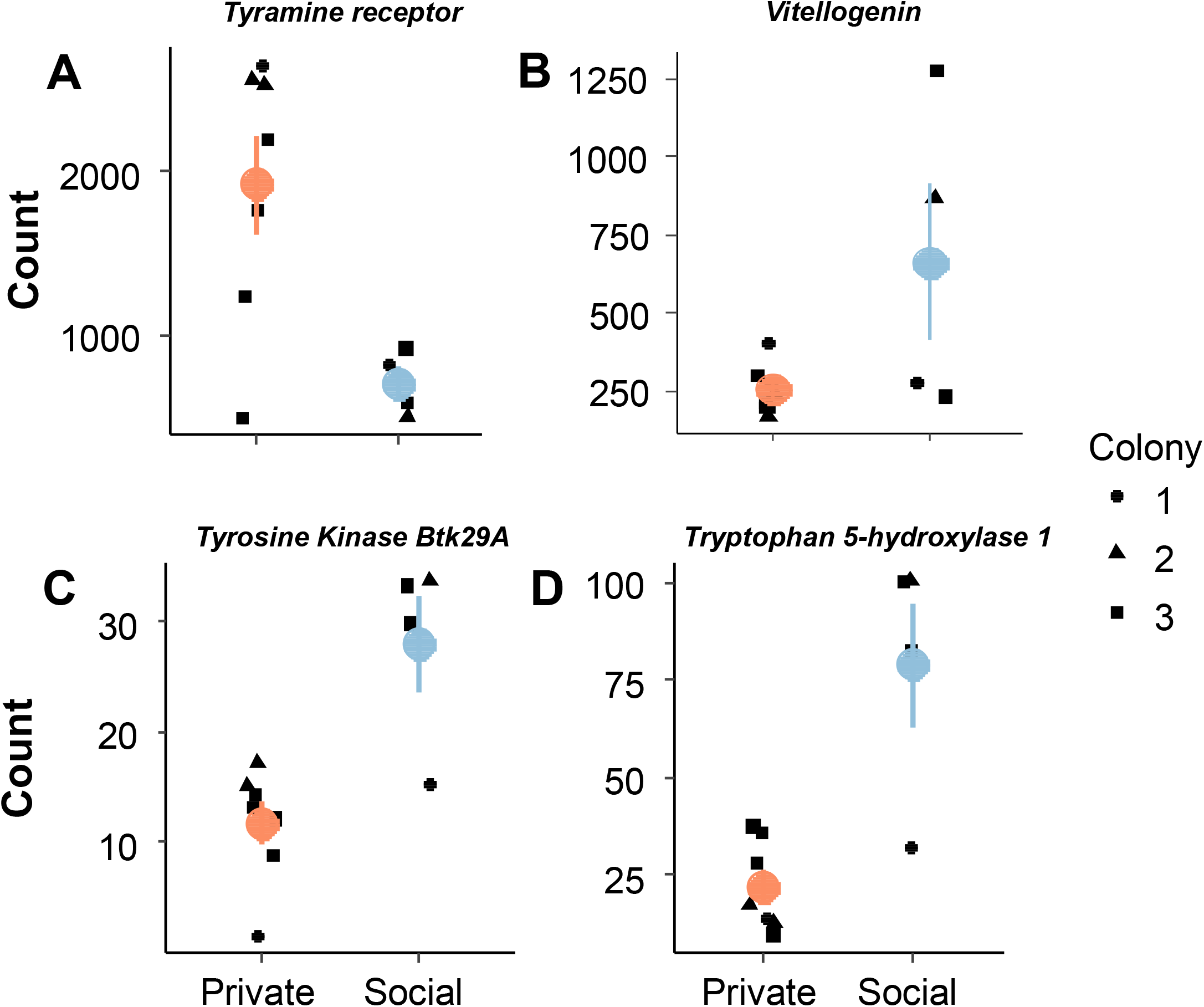
Principal Component Analysis (PCA) plots displaying variance between individual samples based on all genes for each tissue type. Samples are organized by color according to information use strategy (blue = social, red = private) and shapes by colony ID (circles = colony 1, triangles = colony 2, squares = colony 3).

### Sample Collection

On the test day, the day after the 60-minute odor training, 2M sucrose solution with the same scent as used during training was offered only at the dance feeder location, while the training feeder was empty. The sucrose concentration at the dance feeder was high to induce the collecting foragers to perform waggle dances. T.F. trained workers could then decide whether to use social information by following the waggle dances performed by the returning dancers (fly to the D.F.) or disregard the dance vector information and use private information (return to the T.F.). The arrival time and capture time of each individual bee was recorded. Dance and dance following behavior were recorded in the observation colony using a high definition camera to quantify dance following behavior by T.F. foragers. Workers trained to the T.F. that arrived at the D.F location were collected in Eppendorf tubes and immediately preserved in liquid nitrogen; these workers were the social information users. Workers trained to the T.F. feeder that arrived at the T.F. feeder location were collected at a similar time; these workers were the private information users.

### Video Analysis

Videos were analyzed using VLC Media Player. Dances and dance following behaviors were analyzed frame by frame. A worker was only counted as following a dance when she was within one antennal length of a marked dancer during the waggle run phase (Grüter et al., 2013; Linn et al., 2020), which is the component of the waggle dance that encodes the vector information (von Frisch, 1967). We compared the dance following behavior of private and social information users with linear mixed-effects models (LME and GLMM’s). The nlme-package and linear mixed-effects models (LMEs) were used when the response variable was normally distributed (waggles per dance followed). The lme4-package and generalized linear mixed-effects models (GLMMs) were used when the response variable had a Poisson distribution (total number of dances followed) (Zuur et al., 2009). Colony-ID and year (2016 and 2018) were included as hierarchically nested random effects to account for their effects (Zuur et al., 2009).

### Brain Dissection and RNA Extraction

In 2016, we dissected the calyxes of the mushroom bodies and antennal lobes from 14 workers (7 social information users and 7 private information users, 2-3 per colony and type). We confirmed that all social information users followed dances extensively. In 2018 we dissected central brains and subesophageal ganglions from 16 workers (8 social information users and 8 private information users, 2-3 per colony and type), and the antennae from 11 different workers (1-4 per colony and type) (see Fig. 1 in Sen Sarma et al., 2009 for a schematic representation of the brain areas and cut-off lines). The additional handling of the samples after being flash frozen in liquid nitrogen caused the antennae of some samples to be brittle and easily break apart. Different workers were used to ensure that whole antennae could be used for equal extraction of RNA from all samples.

Heads from individual workers were cut from the body and fixed on melted dental wax in a pre-chilled petri dish over ice. The antennae were cut off and stored in 100 mL of TRIzol™ (Invitrogen, USA). Incisions were made at the antennal base, around the eyes, through the compound eye, and the ocellus. The cuticles, glands, retina and tissue around the brain were removed and the exposed tissues of the head were submerged with cooled bee saline (154 mM NaCl, 2 mM NaH__2__PO__4__, 5.5 mM Na__2__HPO__4__, pH 7.2). Subesophageal ganglion and central brain (which included the mushroom body peduncles, the bundled axons from the Kenyon Cells in the calyces), were removed by cutting off optic lobes, antennal lobes, and mushroom body calyces. All tissues called “mushroom body” refer to mushroom body calyces as it is extremely difficult to remove mushroom body peduncles. The calyces contain the intrinsic Kenyon cells, where a large part of mushroom body transcription takes place and, therefore, the calyces are often used to study mushroom body gene expression (Sarma et al., 2009; Reim & Scheiner, 2014; Humphries et al., 2003). Furthermore, the tissue called “central brain” refers to a brain region that also includes the mushroom body peduncles and putative differences in expression in this tissue should be interpreted carefully because of the different functions of these tissues. Each dissection was completed in less than 5 minutes to prevent RNA degradation. Brain parts were stored in 100 mL of Trizol™ (Invitrogen, USA) in −80° for later RNA extraction using RNAeasy Mini Extraction Kit™ (Qiagen, Germany) according to the manufacturers’ protocol.

### Transcriptome Analysis

For sequencing, aliquots of RNA from private and social information users were sent to Beijing Genomics Institute (BGI) for library construction and sequencing. In 2016, Hiseq 4000 was used to sequence 100 base pair (bp) paired-end reads, obtaining 40 Mio clean reads per sample. The total sample size was 28. In 2018, BGISeq was used to sequence 100 base pair (bp) paired-end reads, obtaining 70 Mio clean reads per sample. The sequencing failed for 1 sample and 1 sample was damaged during the travel (Eppendorf tube burst), decreasing our total sample size to 41. Raw reads were quality checked using *FastQC* v.0.11.8 (Andrews et al., 2010) followed by Illumina adapter removal using *Trimmomati*c v.0.38. (Bolger et al., 2014). Clean reads were aligned using *HiSat2* v.2.1.0 (Kim et al., 2017) to the honey bee genome HvA3.1 as a reference (Wallberg et al., 2019). To count how many aligned reads mapped to genes, we used *HtSeq* v.0.11.2 (Anders et al., 2015) to generate count tables. Count tables for each part were analyzed separately for gene expression differences between social and private information users using the R package *DESeq2* v.1.24.0 (Love et al., 2014). Before the analysis, an additional filtering step was added to ensure that only genes with counts of at least 10 reads in at least 6 samples (n-1 of the smallest sample size) were used in the gene expression analysis. Information strategies were compared using the likelihood ratio test (LRT) approach whereby a full model with information type (SI or PI) and colony-ID as fixed factors is compared with a reduced model containing only colony-ID, taking into consideration colony effects. Genes were considered differentially expressed if the false discovery rate (FDR) corrected p-value was < 0.05. To ensure that the number of DEGs calculated by *DESeq2* were not due to chance and to account for the uneven number of samples across bee types and colonies for the antennae, we additionally performed permutations by switching samples from opposite information user groups while maintaining colony structure (see methods in Libbrecht et al., 2016). For example, a sample from the same colony was switched for a different information user group and the number and distribution of DEGs was compared to those calculated from our model in *DESeq2*. We performed 28 permutations (14 times switching two samples for each group and 14 times switching three samples for each group) and recorded the number of DEGs in each permutation. We then compared this number to the numbers for all possible combinations of our samples to assess the number of DEGs that could be expected by chance.

We used the R package *DEGreport* v.1.20.0 (Pantano, 2019) to visualize any patterns for all genes going into the analyses and to identify clustering patterns across social and private information users by using the *rlog* function of *DESeq2* to generate normalized count data and the default settings. PCAs (principal components analysis) based on all genes were performed for all tissues to visualize variation between samples. All analysis were performed in R v.3.5.0 (R Developmental Core Team, 2019).

### Gene Ontology Enrichment

DEGs were loaded in a BLAST search on the NCBI database against the honey bee genome HvA3.1 to find gene annotations. To further obtain information about Gene Ontology (GO) (Ashburner et al., 2000) and KEGG pathway (Ogata et al., 1999) enrichment we used InterProScan v.5.36-75.0 (Jones et al., 2014) on the protein sequences. The R package *topGo* v.2.36.0 (Alexa & Rahnenfuhrer, 2016) was used to perform an enrichment analysis of GO terms and a Fisher’s exact test was performed on the list of biological processes.

## Results

### Dance following of social and private information users

Dance following behavior was analyzed by combining data collected from video analysis for both years. Using a linear mixed-effects model, we found that SI bees followed fewer dances than PI bees during the testing period (5 ± 0.7559 vs. 7.091 ± 1.546 dances) (LME: t-value = 2.218; p = 0.0396). However, SI bees followed dances for longer (more waggle runs per dance) than PI bees (27.214 ± 4.089 vs. 30.818 ± 6.5) (GLMM: z-value = −2.122; p = 0.0338).

### Gene Expression Analysis

The likelihood ratio test (LRT) comparison of information use strategies revealed no differences in gene expression between the two information user groups in the central brain, antennal lobes, and subesophageal ganglion (Fig. 1). There was only one differentially expressed gene between social and private information users from our mushroom body calyxes’ samples, which encodes for an uncharacterized protein (p = 0.026, gene ID: rna-XR_003305479.1). However, there were 413 differentially expressed genes in the antennae, 318 were higher expressed in social information users and 95 were higher expressed in private information users. To confirm these substantial differences in gene expression in the antennae, we used permutations of samples to assess how this affects the number of DEGs in the antennae. The permutations showed that only very few DEGs were found when 2-3 samples were swapped between the SI and PI groups within their respective colonies (colony ID as fixed factor: 11.89 ± 31.87, N = 28; colony ID not included: 3.25 ± 7.01, N = 28) (Fig. S1). This confirms that the substantial differences in gene expression in the antennae are linked to whether bees belonged to the SI or the PI group. PCA plots used transformed data of all genes to further explore whether there is a clustering of samples based on information use strategies and colony. While a clustering pattern based on information use and colony can be seen for the antennae (Fig. 1), the other tissues showed no clear clustering according to information use.

Exploring the list of DEGs in the antennae revealed that numerous odorant binding and chemosensory proteins differed in their expression in social and private information users. Specifically, we detected five genes for odorant or chemical perception among the upregulated genes in private information users (*odorant binding protein 5,11, 19,7 and chemosensory protein 1*) and two among the upregulated genes in social information users (*odorant binding protein 7 and chemosensory protein 2*) (Fig. 2). Several genes involved in biogenic amine production or signaling were also differentially expressed. Social information users had a higher expression of *tyrosine kinase Btk29A; dopamine N-acetyltransferase, tryptophan 5-hydroxylase 1,* which are involved in the production of dopamine or serotonin (Vavricka et al., 2014; Coleman et al., 2005; Sasaki et al., 2012), while private information users had a higher expression of one gene *tyramine receptor, transcript variant X1*, which is associated with biogenic amine signaling (Mustard et al., 2005; Blenau et al., 2000) (Fig. 3). Social information users also had higher expression of the egg yolk precursor protein *vitellogenin*, a gene that is upregulated in nurses and downregulated in foragers fat bodies and brain (Amdam et al., 2002; Nunes et al., 2013) (Fig. 3).

**Figure 2:**
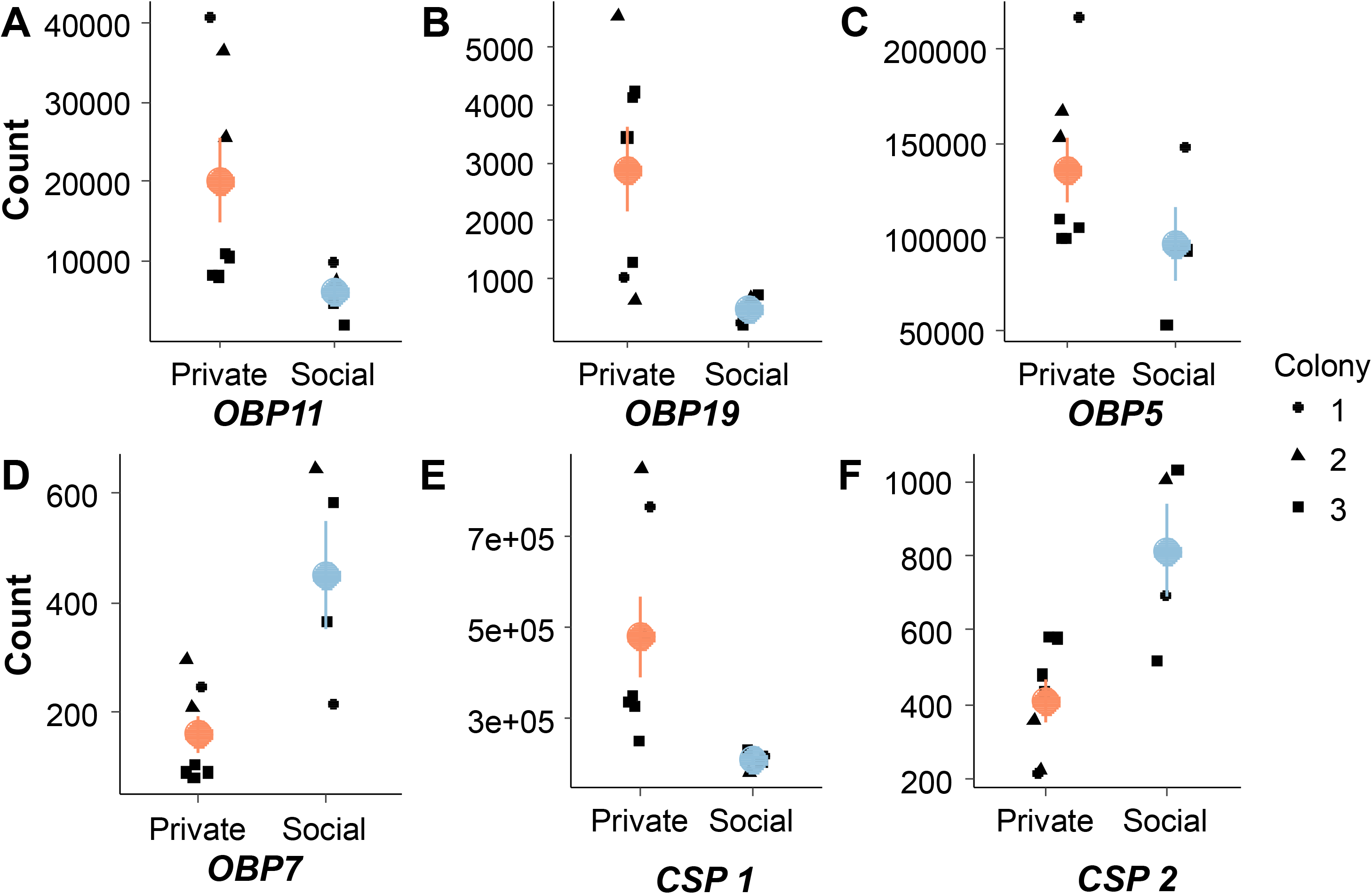
Plots of individual odorant binding protein (OBP) and chemosensory protein (CSP) genes. Black dots show counts for individual samples and shapes correspond to the colony ID (circle = colony1, triangle = colony 2, square = colony 3). Colored circles are representative of the average for the respective information strategy (red = private, blue = social) with confidence intervals. A) OBP11 (p<0.001) B) OBP19 (p=0.001) C) OBP5 (p=0.03) D) OBP7 (p<0.001) E) CSP1 (p=0.009) F) CSP2 (p=0.007). P-values shown are after FDR correction.

**Figure 3:**
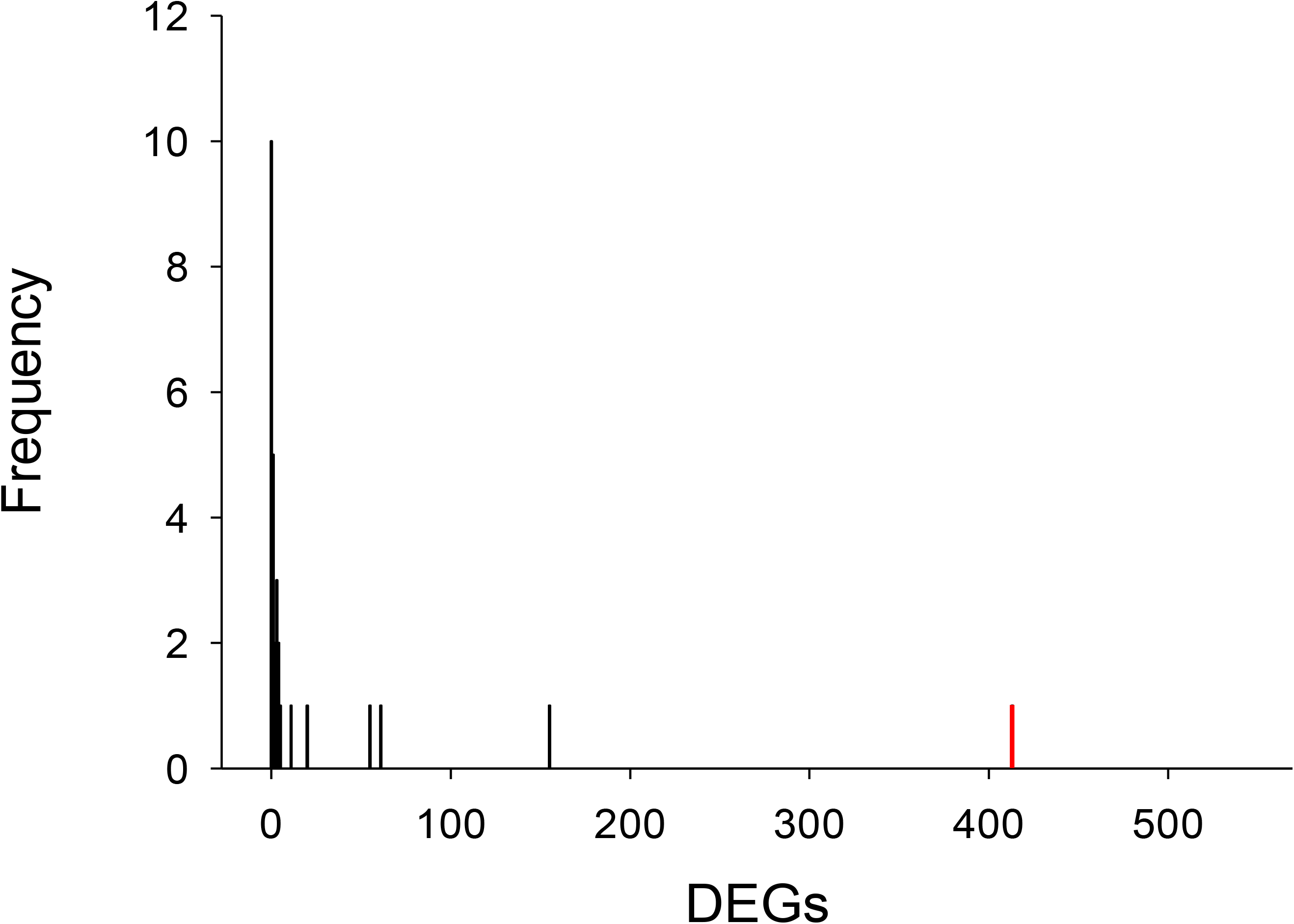
Plots of individual genes associated with biogenic amine production and reproduction. Black dots show counts for individual samples and shapes correspond to the colony ID (circle = colony1, triangle = colony 2, square = colony 3). Colored circles are representative of the average for the respective information strategy (red = private, blue = social) with confidence intervals. A) Tyramine receptor (p=0.018) B) Vitellogenin (p = 0.045) C) Tyrosine Kinase Btk29A (p=0.006) D) Tryptophan 5-hydroxylase 1 (p<0.001). p-values shown are after FDR correction.

### Gene function and enrichment analysis

Separate GO enrichment analyses of only upregulated genes for each information strategy showed a small number of enriched functions: 9 biological processes enriched in social information users connected mainly on *carbohydrate (10 genes) and lipid (7) metabolic process* and 18 enriched biological processes in private information users focused on *oxidation-reduction process (7) and protein catabolic process (11)*.

## Discussion

Information and its use in animals is an important topic in behavior, ecology, and evolution because information is a critical currency that allows animals to make adaptive decisions in a given situation (e.g. Danchin et al., 2004; Dall et al., 2005; Rieucau & Giraldau, 2011; Hoppitt & Laland, 2013). The decision of when to utilize social versus private information to best exploit potential opportunities while avoiding costs is crucial for success and has been studied within a variety of both social and non-social animals (e.g. Bonnie & Earley, 2007; Weimerskirch et al., 2010; Grüter & Ratnieks, 2011; Wray et al., 2011; Taborsky & Oliveira, 2012; Haak et al., 2020). However, it is still unclear if and how molecular and neurosensory factors determine an individual’s preference for social or private information.

Here we explore whether gene expression differences between honey bee foragers are linked to the use of social or private information about food sources to uncover the potential molecular mechanisms that underlie the decision to decode and use waggle dances in honey bees. Contrary to our prediction, the transcriptomes of all four analyzed brain parts did not differ between bees using these two foraging strategies. Strikingly, however, we found substantial gene expression differences in the antennae. Over 400 genes were differentially expressed between social and private information users, suggesting that the sensory perception of these two forager types differs. This is further supported by expression differences related to odorant binding proteins, chemosensory proteins, and genes associated with biogenic amine production.

The lack of differences in the brain areas was unexpected given that Liang et al. (2012) found extensive differences in whole-brain gene expression between scouts and non-scout foragers (in their study, non-scout foragers could have included both private and social information users). We expected the mushroom bodies to show differences since it has previously been shown that they are involved in multisensory integration, learning, and place memory (e.g. Strausfeld et al., 2009; Collett & Collett, 2018). The antennal lobes are involved in odor recognition and memory through the interconnectivity of neurons with the mushroom body and were thus selected as another area of interest (Boeckh & Tolbert, 1993). The central brain has been suggested as an important area for dance communication (Barron & Plath, 2017), while the subesophageal ganglion plays important roles in reward perception and taste (Galizia et al., 2011). Together, these brain regions were thought to process reward and odor perception which could play an important role in the decision to use dance information. Our study indicates that information use strategies may not primarily depend on integration of information in higher order centers, but that the antennae play a major role in decision-making when facing communication signals.

The 413 differentially expressed genes in the antennae present an array of gene families and functions. Of particular interest are genes coding for odorant binding proteins and those involved in biogenic amine production and signaling due to their potential roles in chemosensory perception. Thus, differences in the perception of chemosensory information cues and signals could result in divergent foraging strategies. While our study cannot disentangle whether gene expression is the cause or the consequence of the information use strategy, they suggest that chemosensory perception by the antennae could be involved in the decision to decode waggle dances and use social information. In many social insects, the antennae play an integral role in social recognition (Ozaki et al., 2005; Sharma et al., 2015; Balbuena & Farina, 2020). Studies in *Oecophylla smaragdina,* for instance, indicated that the density of antennal sensilla is important in regulating behavior, particularly in determining the aggression response behavior to non-nestmates (Gill et al., 2013; Cholé et al., 2019). Similar to other social insects, honey bee foragers first use their antennae to perceive and respond to a variety of chemical signals for navigation (Menzel & Greggers, 2013), efficient nectar/pollen collection (Arenas & Farina, 2012), and dance communication (Thom et al., 2007; Reinhard & Srinivasan, 2009; Gilley et al., 2012).

By transporting odorants, e.g. from antennal sensilla to odorant receptors, odorant binding proteins (OBPs) play important roles for olfactory sensitivity (Leal, 2013). They are hypothesized to be important in insect communication (Pelosi et al., 2005), including in the honey bee which use highly complex odors and pheromones to regulate their social activities (Farina et al., 2012; Baracchi et al., 2020). Of the 21 OBPs found in the honey bee, only 9 are exclusively expressed in the antennae. The remaining OBPs are active throughout the honey bee body or specific non-olfactory tissues (Forêt & Maleszka, 2006). Our analysis revealed that workers which rely on private information in the form of spatial memory show higher expression of four odorant binding proteins (*obp5*, *obp11*, *obp19*, and *obp7*), whereas workers that rely on socially acquired information upregulate one (*obp7*). Thus, ~25% of all OBPs found in honey bees were differentially expressed. Of the OBPs that were upregulated in private information users, *obp5* and *obp11* have been previously shown to be exclusively expressed in the antennae and suggest a chemosensory function (Forêt & Maleszka, 2006). Interestingly, *obp11* is mainly expressed in a rare type of antennal sensilla found only in female honey bees, the *sensilla basiconica*, and is likely to facilitate the function of these sensilla (Kucharski et al., 2016). While the ligand of *obp11* remains unknown, there is evidence that the *sensilla basiconica* play important roles in the perception of cuticular hydrocarbons (CHCs) in ants (Sharma et al., 2015) and may play a similar role in honey bees (Kucharski et al., 2016). This is remarkable because CHCs emitted by dancing bees are known to trigger the use of private information in honey bees (Thom et al., 2007; Gilley et al., 2012). This raises the possibility that a higher expression of *obp11* increases the sensitivity of bees towards CHCs emitted by waggle dancers, thereby triggering private information use. The remaining differentially expressed OBPs (*obp19* and *obp7*) have been shown to be ubiquitously expressed, which suggests they may have additional molecular functions which we currently do not know. Overall, these results indicate a difference in perceptional sensitivity where workers which use private information perceive some chemosensory stimuli more or differently than social information users. This could have far reaching consequences for their behavior given the role that odors play in the decision-making and information use of a forager, e.g. in the identification and learning of floral resources or the perception of cuticular hydrocarbons (von Frisch, 1967; Johnson, 1967; Reinhard et al., 2004; Grüter et al., 2008; Gilley et al., 2012).

Chemosensory proteins serve a similar role as OBPs in transporting chemical stimuli through mechanisms that are not yet well understood. These proteins are heavily concentrated in antennal sensilla but are also expressed in non-olfactory tissues (Forêt et al., 2007; Calvello et al., 2005). Of the six chemosensory proteins found in honey bees (McKenzie et al., 2014), two were differentially expressed in social and private information users, chemosensory proteins 1 and 2. Both chemosensory proteins have been shown to be highly expressed in the antennae (Li et al., 2016), which further supports the view that the differences between the information strategies may be rooted in chemoreception.

Biogenic amines have been associated with regulating learning, foraging behavior, and the transition from in-hive tasks to foraging (Lehman et al. 2006). Biogenic amine signaling is known to change with age and tissue location in honey bees (e.g. McQuillan et al., 2012; Perry & Barron, 2013; Reim & Scheiner, 2014; Thamm et al., 2017). Specifically, dopamine, serotonin, octopamine, and tyramine titers in the brain were found to be linked to both task and age (Schulz & Robinson, 1999; Barron et al., 2002; Harris & Woodring, 1991; Kokay & Mercer, 1997). For example, tyramine levels have been linked to novelty seeking in scouting behavior (Cook et al., 2018; Liang et al. 2012), sucrose responsiveness (Scheiner et al., 2002; 2017a; 2017b), and division of labor between nectar and pollen foragers (Hunt et al., 1995; Scheiner et al., 2001). Dopamine has been shown to modulate sucrose responsiveness (Scheiner et al., 2002), learning (Vergoz et al., 2007) and dance following (Linn et al., 2020), whereas serotonin influences foraging activity (Schulz et al., 2003) and regulates feeding in many animals (French et al., 2014; Blundell & Halford, 1998; Voigt & Fink, 2015). Our findings of an upregulation of genes associated with biogenic amine production, raise the possibility that social information users could differ in their sensory perception as well as sucrose response thresholds compared to private information users. It is noteworthy, however, that the differences we found in relation to biogenic amine signaling were not in the brain. Instead, higher expression of several genes associated with dopamine and serotonin production was found in the antennae of social information users. We did not control for foraging age or experience, which have already been shown to affect gene biogenic amine expression (Reim & Scheiner, 2014). However, the lack of differential expression in brain areas suggests that there was no systematic age bias in our samples.

Another interesting differentially expressed gene, *vitellogenin*, is best known as an egg yolk precursor protein for egg laying organisms. Under normal conditions in social insects, the queen is the main reproductive member and therefore produces the highest levels of *vitellogenin*. However, *vitellogenin* serves important roles for other behaviors and functions outside of reproduction (Nelson et al., 2007; Morandin et al., 2014). For example, nurses produce the next highest levels of *vitellogenin* in their hypherengeal glands to fortify brood food with protein (Amdam et al., 2003; Amdam et al., 2009; Wegener & Bienefeld, 2009). A characteristic feature of the transition from nurse to forager is the drop in vitellogenin levels (Amdam et al., 2003; Messan et al., 2018). Our finding is consistent with evidence that biogenic amine levels are linked to *vitellogenin* and foraging behavior (Linn et al., 2020; Koywiwattrakul et al., 2005), where social information have a similar physiological state to nurses.

Intrinsic factors such as genetic differences could also affect the decision to decode waggle dances. Honey bee queens can mate with more than 20 drones (Strassman, 2001), and the patriline composition of our samples is not known. It is well-known that different patrilines can differ in foraging behaviors, such as foraging age (Kolmes et al., 1989). Paternal effects can also impact gustatory responsiveness and learning abilities (Scheiner et al., 1999; 2001; 2005; Behrends et al., 2007; Scheiner & Arnold, 2009). It is unclear whether systematic patriline differences in the composition of PI and SI bees would lead to differential gene expression only in the antennae, but future studies should explore whether bees using private or social information differ in their patrilines.

Overall, our results suggest an important role of the antennae in mediating decision-making and information use. In particular, we suggest a link between chemosensory perception and the reliance on communication in honey bees. Further studies are needed to disentangle the potential effects of genetic differences (i.e. different patrilines), differences in foraging experience, and other factors on gene expression. In addition, we need studies to confirm our hypothesis that SI and PI bees differ in sensory perception such as sucrose response thresholds, odor learning, and electroantennograms.

## Acknowledgements

We are grateful to Semi Brami for help with data collection and Romain Libbrecht and Marah Stoldt for their support in the gene expression analyses. We thank Ricarda Scheiner and Markus Thamm for helpful advice on the brain dissections. We also would like to thank the German Research Foundation (DFG) for a grant to C. G. and S. F. (DFG GR 4986/3-1 and FO 298/27-1). T.P. was supported by a fellowship of the China Scholarship Council (File No. 201606170134). S.M.G. was supported by a grant from the ‘Inneruniversitäre Forschungsförderung’ of the University of Mainz. Our laboratory work was supported by Marion Kever and Steffi Emmling.

## Author Contributions

A.K., T.P., M.L., S.M.G., S.F. and C.G. conceived the study and designed the experiments. A.K., T.P., M.L. and S.M.G. conducted the field experiments. T.P., M.L. and S.M.G. performed all honey bee brain dissections and RNA extractions. T.P and A.K worked together on gene expression analysis. Data analysis was supported by S.F and C.G. A.K. wrote the first draft of the manuscript, all authors contributed to writing the final version.

## Data Accessibility

All of the supplemental material and additional data generated and used throughout this project may be found within the Dryad repository https://doi.org/XX.XXXX/dryad.XXXXXX, which contains the following: dance following information for all samples, lists of differentially expressed genes for all brain tissues, InterProScan annotation of gene lists and GO enrichments from all antenna samples. All sequencing data will be deposited in the Sequencing Read Archive (SRA) of the NCBI upon acceptance.

---

**Supplemental Figure 1:**
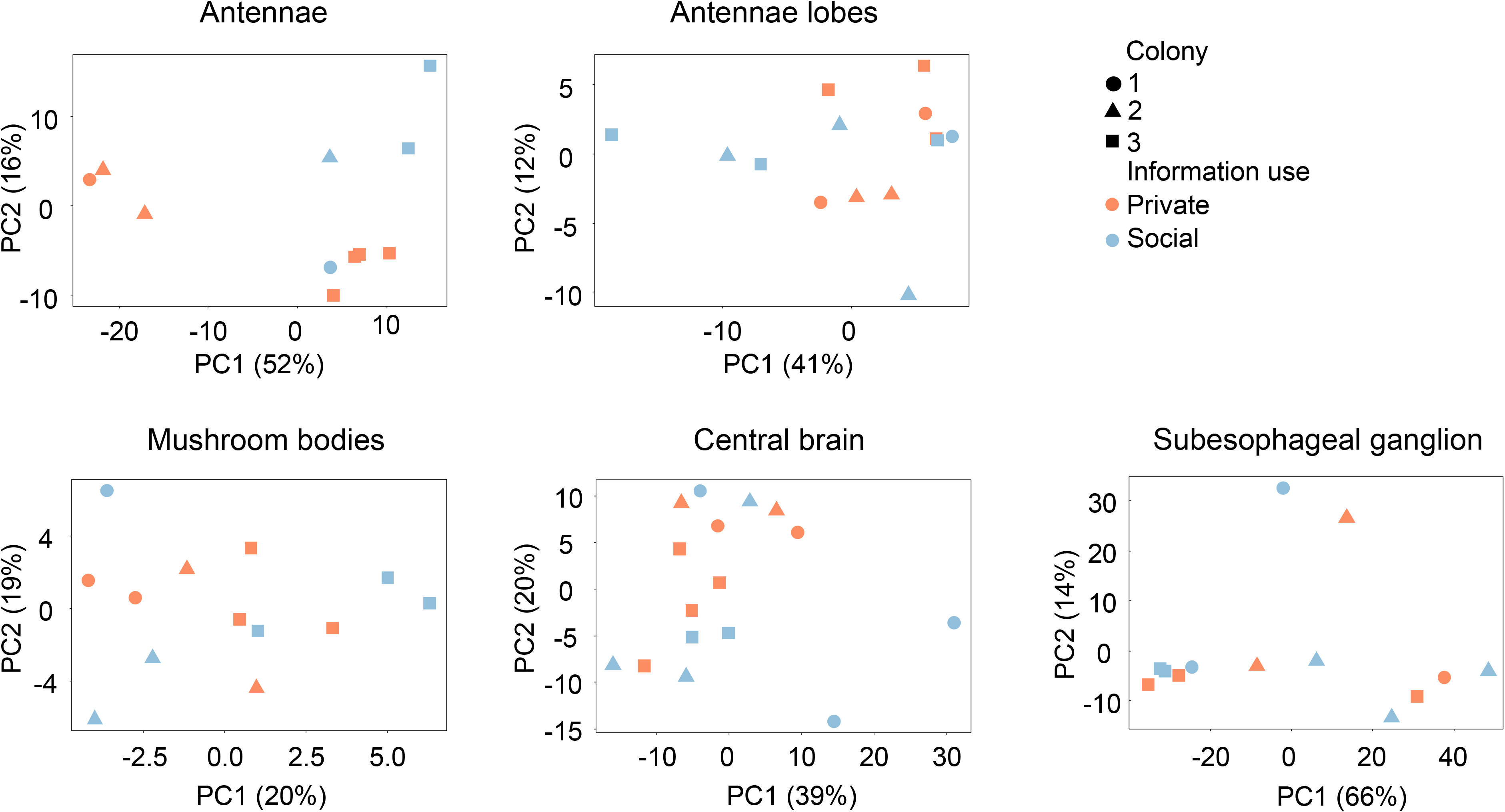
Distribution of differentially expressed genes (DEGs) expected by chance after permutation results (N = 28). The x-axis indicates the number of DEGs and the y-axis indicates the frequency of occurrences. Black bars indicate the results from the permutations, the red bar indicates our results.

